# Scalable cytoarchitectonic characterization of large intact human neocortex samples

**DOI:** 10.1101/274985

**Authors:** Sven Hildebrand, Anna Schueth, Andreas Herrler, Ralf Galuske, Alard Roebroeck

## Abstract

We describe MASH (Multiscale Architectonic Staining of Human cortex): a simple, fast and low-cost cytoarchitectonic labeling and optical clearing approach for human cortex samples, which can be applied to large formalin fixed adult brain samples. A suite of small-molecule fluorescent nuclear and cytoplasmic dyes in combination with new refractive index matching solutions allows deep volume imaging. This enables highly scalable human neocortical cytoarchitecture characterization with a large 3D scope.

---

Optical volume imaging of human cerebral cortex is challenging at the cellular scale owing to the large size of the human brain and the 3-dimensional geometry of the cortex. The 2-4mm thick cortical sheet is highly curved and packed with billions of neurons, organized in layers each hundreds of micrometers thick. Traditionally, studies on human cortical cytoarchitecture have been performed on sections with a thickness of less than 100µm. However, thin sections have no clear geometric relation to the curved cortical sheet, mostly slicing it non-orthogonal to the layer organization. Due to shape distortions and tearing inherent to the sectioning process, serial sections are extremely difficult to align post-hoc into a volume dataset. Although a recent surge of optical clearing techniques has transformed microscopic 3D imaging of small transgenic or antibody stained rodent brains^1–6^, translation of these techniques to the much larger adult human brain has remained a challenge. More specifically, volume imaging and cytoarchitectonic characterization of large, adult formalin fixed brain samples, scalable in terms of time and cost to thousands of cubic millimeters in order to cover a significant portion of a human cortical area, has so far remained out of reach.

Here, we report MASH (Multiscale Architectonic Staining of Human cortex): a novel scalable nuclear and cytoplasmic labeling and optical clearing approach. MASH is suitable for 4-5mm thick archival (i.e. formalin fixed and long-term stored) adult human cortex samples and enables high-throughput deep 3D optical imaging. MASH consists of two innovations: 1) a set of low-cost small-molecule fluorescent dyes and cleared tissue cytoarchitecture labelling protocols (MASH dye protocols) and 2) a set of adjustable refractive index matching solutions (MASH RIMS) which enable the use of the MASH dye protocols in cleared tissue. For the MASH dye protocols, we identified four small organic compounds: acridine orange (AO),methylene blue (MB), methyl green (MG) and neutral red (NR), previously used as cytoplasmic and nuclear labels in traditional light microscopy studies e.g.^7^, ^8^. We developed adapted protocols for their use as fluorescent labels in large cleared human brain specimen: MASH-AO (green spectrum cell-body label), MASH-NR (red spectrum cell-body label), MASH-MB (far-red spectrum cell-body label) and MASH-MG (far-red spectrum cell-nucleus label). In order to apply the MASH dye protocols in a wide range of human cortex samples, the clearing process must be: 1) potent enough to clear highly myelinated adult human brain tissue up to 4-5 mm thickness within reasonably short time, 2) compatible with MASH dye protocols and, ideally, 3) applicable to archival samples available from brain banks and other academic and clinical tissue storing facilities. The DISCO family of clearing protocols ^3^, ^5^, ^9^ have short clearing times and they have been applied to freshly frozen ^10^ and 1 mm thick formalin fixed ^9^ human brain tissue. A serious challenge in these protocols is posed by the refractive index matching solutions (RIMS): di-benzyl ether (DBE) or mixtures of benzyl alcohol (BA), benzyl benzoate (BB) and diphenyl ether (DPE). They are not compatible with imaging of MASH dyes, either due to their photo bleaching properties or their high melting point. Moreover, their corrosiveness limits the microscope setups which can be used due to the detrimental effects on many microscope objectives. Therefore, we developed an adapted DISCO approach replacing the RIMS with two alternative new MASH RIMS. We identified the essential oil *trans*-cinnamaldehyde (CA) as a high refractive index (RI) medium, mixable with Thiodiethanol (TDE) or wintergreen oil (WGO) to create TDE/CA and WGO/CA RIMS. The MASH RIMS have the properties of low fluorescence quenching, low melting point, low corrosiveness, and are fully compatible with the MASH dye protocols.

We demonstrated that MASH can clear and label thick (4-5mm) archival adult cortex samples and that cytoarchitecture can be imaged at a variety of wavelengths, depths and magnifications (**Fig. 1**). Volume imaging can be performed in the green spectrum (MASH-AO, **Fig. 1a,b,d**; **Supplementary Video 1**), red spectrum (MASH-NR, **Fig. 1c,g,i-k**) and far-red spectrum (MASH-MB, MASH-MG, **Fig. 1e,f**; **Supplementary Video 5**). This imaging can be performed at sub-micron resolutions using Two-photon microscopy (TPM) to delineate single neuron cell-body morphology and cortical layer borders (**Fig. 1a-d**). Likewise, imaging can be performed over larger fields-of-view up to the entire cortical sheet with light sheet fluorescence microscopy (LSFM; **Fig. 1e,f**). Clearing and staining can be performed on 5mm thick samples in a matter of 10 days and result in imaging of cytoarchitecture with low background and high signal over the full imaging depth (**Fig. 1g-k**). MASH cell body labelling is well co-localized with standard Nissl stain cresyl violet (CV) in thin sections showing its labeling specificity and suitability for cytoarchitecture characterization (**Fig. 1l-n**). Labeling specificity was further validated by verifying co-localization with the DAPI nuclear stain and bright-field cytoplasmic stains (**Supplementary Fig. 1,2**).

**Figure 1:**
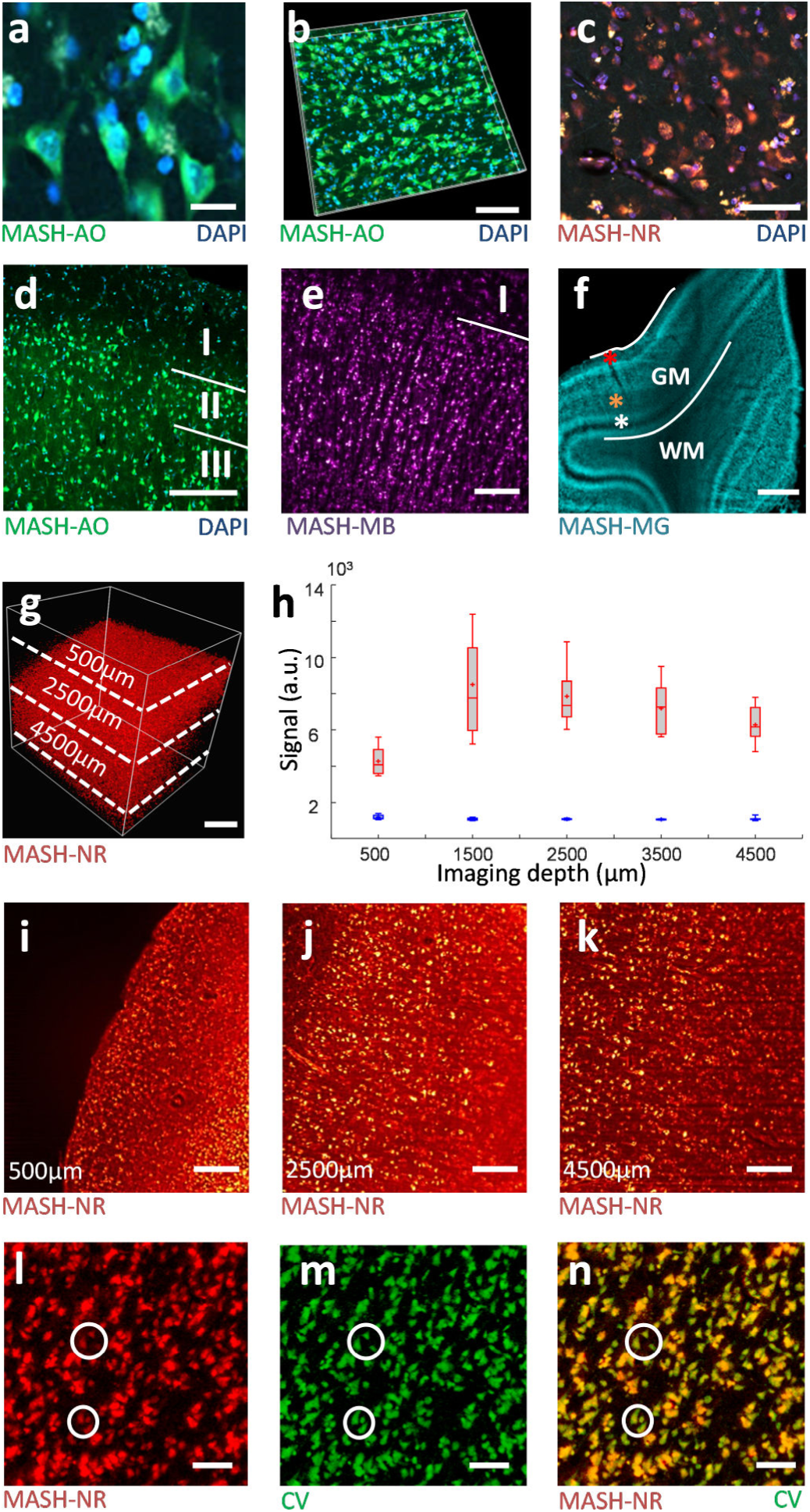
MASH labels human cortical cytoarchitecture in cleared formalin fixed tissue imaged at high resolution and depth. (A) Two-photon microscopy (TPM) image of a cleared human neocortex sample stained with MASH-AO for neuronal cell-bodies (green), counterstained with DAPI (blue) for cell-nuclei. (b) 3D rendering of the corresponding imaging stack. (c) TPM image of a cleared sample stained with MASH-NR for neuronal somata (red) and DAPI (blue). (d) TPM image of MASH-AO and DAPI stain showing the layer I/II and II/III borders (e) light sheet fluorescence microscopy (LSFM) of MASH-MB showing the layer I/II border (f) low-magnification LSFM imaging of MASH-MG stain showing the entire gray matter (GM), the white matter (WM) transition and cortical layer contrast (red asterisk: layer I; orange asterisk: stripe of Gennari, layer IVb; white asterisk: inner stripe of Baillarger, layer V). (g) 3D rendering of 4.5mm deep (referenced to tissue surface) LSFM imaging stack of MASH-NR stained tissue (h) Mean, median and spread (n=15) for neuronal cell body signal (red boxplots) and non-neuronal tissue background signal (blue boxplots) at several depths in g. (i-j-k) Images indicated in g at 500µm, 2500µm and 4500µm imaging depth. (l-n) Validation by bright-field cresyl violet stain (CV) shows co-localization of somata in a 50 µm thick section of human cortex. (l) the section stained with MASH-NR (red) and (m) with CV (green). (n) Overlay of l + m, circles: corresponding locations. Scalebars a:20 µm; b,c:50 µm; d,l,m,n: 100 µm; e,i,j,k: 200 µm; f,g: 1mm

**Figure 2:**
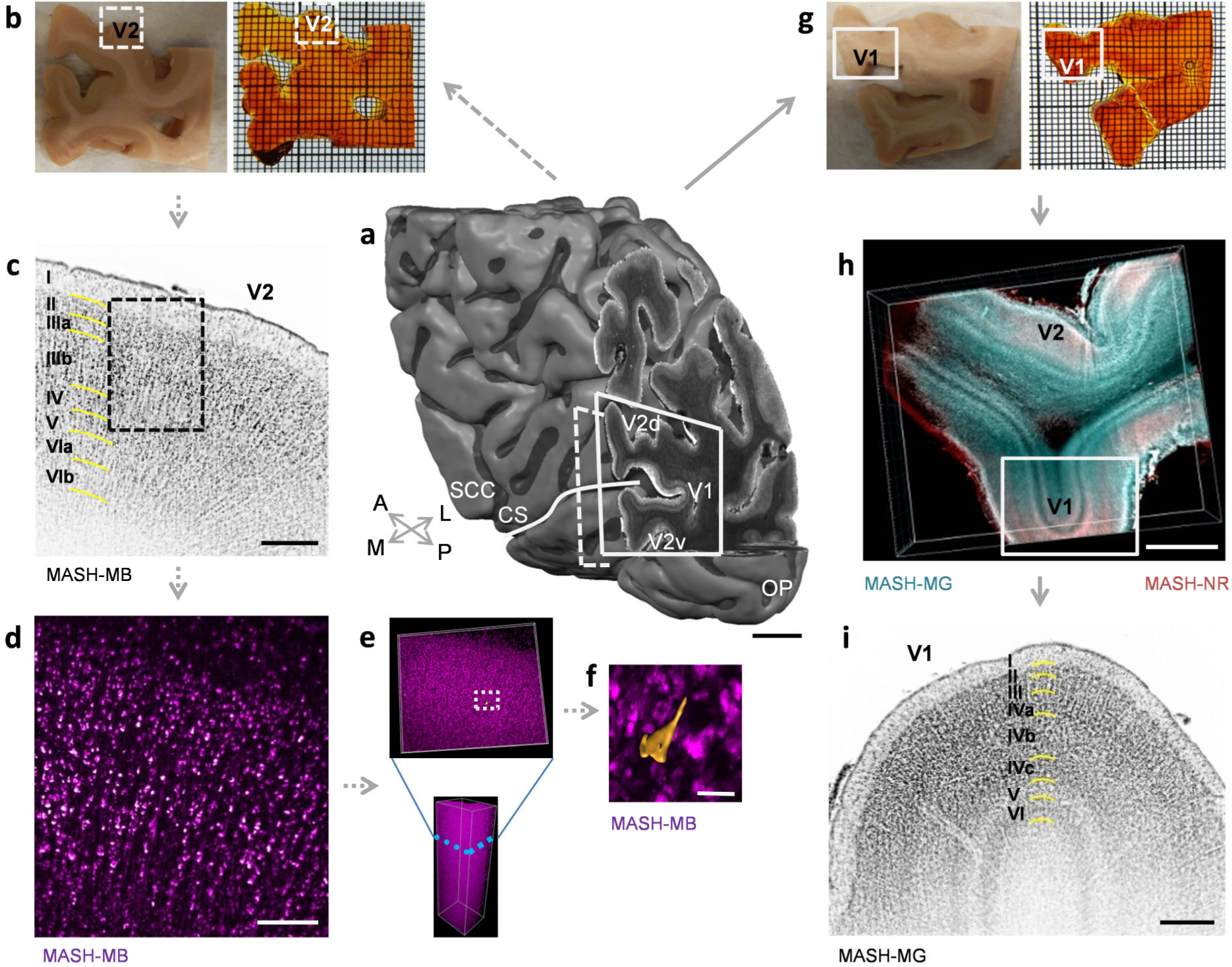
MASH allows characterisation of human cortical cytoarchitecture in large formalin fixed samples over a large range of scales. (A) 3D MRI reconstruction of the human occipital lobe sample with primary and secondary visual cortex around the calcarine sulcus. V1: primary visual cortex; V2: secondary visual cortex; V2v: ventral V2; V2d: dorsal V2; CS: Calcarine Sulcus (white line); SCC: Splenium of the Corpus Callosum; OP: Occipital Pole; A: anterior; P: posterior; M: medial; L: lateral. (b) The anterior 5mm thick sample (dashed line in a), before (left) and after staining and clearing (right). (c) LSFM imaging of the MASH-MB stain in V2 (inverted greyscalemap) in the dashed box in b, with cytoarchitectonic layering characterisation (left). (d) Higher magnification LSFM of the dashed box in c. (e) the entire 3D LSFM stack at the position of d with magnification of a sub-stack at the blue dashed line (f) a 3D surface reconstruction of a pyramidal neuron cell body from LSFM data at the dashed box in e, imaging depth 1066µm. (g) The posterior 5mm thick sample (solid line in a), before (left) and after staining and clearing (right). (h) LSFM imaging of dual MASH-NR soma staining and MASH-MG nucleus staining in V1 and V2 in the dashed box in g. (i) higher magnification LSFM imaging of the MASH-MG channel in the solid box in h (inverted orientation), with cytoarchitectonic characterisation of V1 cortical layering (middle). Thin grid (b,g): 1mm. Scalebars: a: 10mm; h: 2mm; c,i: 500µm; d: 200µm; f= 50µm.

We demonstrated the efficacy of MASH in high-throughput 3D LSFM characterization of human cortical cytoarchitecture by applying it to large (∼40x30x5mm) archival samples surrounding the calcarine sulcus (**Fig. 2**). The samples were large enough to be easily localized in a magnetic resonance imaging (MRI) reconstruction of the entire host occipital lobe and to contain parts of both primary (V1) and secondary (V2) visual cortical areas (**Fig. 2a**). Clearing was highly effective, rendering the entire 5mm thick samples transparent with a slight amber-like tint (**Fig. 2b,g**). Labeling could be achieved over the entire depth of the samples with the far-red MASH-MB cell-body stain for the anterior sample (**Fig. 2b-f**) and a two-color labeling with the red MASH-NR cell-body stain and the far-red MASH-MG nuclear stain for the posterior sample (**Fig. 2g-i**). Multi-spectral LSFM imaging of layered cytoarchitecture could be performed at low magnification over 10-12mm long stretches of V1 and V2 cortical sheet (**Fig. 2h**). Higher magnification LSFM produced mesoscale imaging volumes with both the resolution to resolve single neurons and the field-of-view to contain all layers of the cortical sheet (**Fig 2c,i**; **Supplementary Video 4**). Planes from these volumes allowed for classification of cortical layering and sub-layering, corresponding to a histological reference atlas ^11^. They displayed distinctive cytoarchitectonic features such as the cell-poor layers IVb and V in V1 (**Fig. 2i**) and the large pyramidal cells in layer IIIb of V2 (**Fig. 2c**). High magnification light sheet imaging (**Fig. 2d**; **Supplementary Video 2,3**) showed soma morphology features of individual neurons in the context of a deep 3D imaging stack (**Fig. 2e**) with sufficient resolution for a surface reconstruction of a pyramidal neuron cell body (**Fig. 2f**).

Applied together, the MASH dye protocols and MASH RIMS allow clearing and labeling of thick adult human brain samples and deep volume imaging of cytoarchitecture. The entire MASH protocol for clearing and labeling of 5 mm thick samples takes approximately 10 days. MASH dye solution costs are low, less than 1$ per sample. This makes it easily scalable to the investigation of the human brain and feasible to apply in standard lab environments. Moreover, MASH is capable of clearing and labeling adult human archival brain samples, even after prolonged storage in formalin (current samples had been fixed for 14 to 30 months), making it applicable to tissue stored in brain banks rather than being limited to fresh or freshly frozen tissue. The low corrosive MASH RIMS can be adapted to various RIs which potentially allows their application in a wider range of optical clearing protocols. Furthermore, samples can be stored long-term in MASH RIMS before imaging because they maintain transparency and fluorescence and do not solidify at low temperatures (2°C to 7°C). In the context of smaller and thinner samples, where antibody labels are effective, or in transgenic animal experiments, the MASH labels can serve as counterstains providing highly useful contextual information in complementary spectral bands. Furthermore, they could be combined with deeply penetrating small-molecule pathology labels e.g. ^10^, to investigate human disease pathology in its full cytoarchitectonic context. We demonstrated LSFM volume microscopy imaging over a variety of scales with a commercially available light sheet system, both at high resolution and at large field-of-view. MASH can be combined with advances of LSFM technology, such as the dual inverted selective plane illumination microscopy (diSPIM) system geometry ^12^, and large-volume image stitching ^13^. This opens new possibilities of cellular level volume imaging of entire human cortical subsystems. Combination with other imaging modalities, such as MRI, can additionally provide correlative multi-modal multi-scale data on cytoarchitecture in the human brain.

## METHODS

### Human brain tissue

Brain tissue samples were taken from 3 different human body donors (no known neuropathological diseases) of the body donation program of the Department of Anatomy and Embryology, Maastricht University. The tissue donors gave their informed and written consent to the donation of their body for teaching and research purposes as regulated by the Dutch law for the use of human remains for scientific research and education (“Wet op de Lijkbezorging”). Accordingly, a handwritten and signed codicil from the donor posed when still alive and well, is kept at the Department of Anatomy and Embryology Faculty of Health, Medicine and Life Sciences, Maastricht University, Maastricht, The Netherlands.

Brains were first fixed *in situ* by full body perfusion via the femoral artery. Under a pressure of 0.2 bar the body was perfused by 10 l fixation fluid (1.8 vol% formaldehyde, 20% ethanol, 8.4% glycerine in water) within 1.5-2 hours. Thereafter the body was preserved at least 4 weeks for post-fixation submersed in the same fluid. Subsequently, brains were recovered by calvarian dissection and stored in 4% paraformaldehyde in 0.1 M phosphate buffered saline (PBS) for 14-30 months.

For MASH clearing and whole-mount labelling procedures, tissue of an occipital lobe (subject 1, fixation time 14 months), occipital and parietal samples (subject 2, fixation time 30 months) and a temporal lobe sample (subject 3, fixation time 3 months) were used (**Supplementary Table 1**). All tissue was manually blocked with anatomical trimming blades, then cut into 2 to 5 mm thick slices in coronal orientation and immediately processed.

### MASH protocol for clearing and labelling of human brain samples

For clearing of formalin fixed adult brain tissue an adaptation of the iDISCO+ method^14^ was used. The incubation times were adjusted for better clearing of formalin fixed adult human brain tissue and dibenzyl ether (DBE) was replaced by two new MASH refractive index matching solutions (RIMS) adjusted to an RI of 1.56, as described below. The MASH protocol consists of the following steps: 1) sample pre-treatment with methanol and bleaching, 2) labelling with MASH dye protocols and 3) Clearing and refractive index matching with MASH RIMS.

### Sample pre-treatment with methanol and bleaching

For the pre-treatment, samples were dehydrated in ascending concentrations of methanol (VWR International, LLC, 20846.326) in distilled water: 1 h each in 20 %, 40 %, 60 %, 80 %, and twice in 100 %. Samples were cooled down to 4 °C during the second incubation in 100 % methanol. This was followed by bleaching with freshly prepared 5 % H_2_O_2_ (1 volume of 30 % H_2_O_2_ for 5 volumes of methanol, ice cold) at 4 °C overnight under shaking. After bleaching and re-equilibration to room temperature (RT) the tissue was rehydrated as follows: incubation for 1 h in 80 %, 60 %, 40 % and 20 % methanol in distilled water and twice in in 0.1 M PBS/0.2% Triton X-100 (VWR International, LLC, 28817.295) respectively. After this, labelling was performed as described below. For all steps, incubation was done in 6 well cell culture plates (Corning Inc., 3516) in a volume of 5 ml/well at RT, unless mentioned otherwise.

### Labeling procedure with MASH dye protocols

Four dyes were identified for cell body (cytoplasm and nucleus) staining or nuclear staining. These dyes, used variously before in animal studies as bright-field or fluorescent stain on histological sections, were investigated as to their suitability as labels of large, cleared adult human specimen for deep fluorescent light microscopy imaging. The four dyes acridine orange (AO), methylene blue (MB), methyl green (MG) and neutral red (NR), see **Supplementary Table 4**, are referred to as MASH-AO, MASH-MB, MASH-MG and MASH-NR respectively in the context of human cleared tissue labelling protocols, to distinguish them clearly from other prior uses. All MASH labels are small organic compounds with low molecular weight and fluorescent properties which have hitherto not been applied for labelling in thick, cleared human tissue. AO and NR have been variously described before as bright-field or fluorescent Nissl stains on standard histological sections^15^. MG was recently described as an effective, low-cost DNA stain, and applied to chick embryo cryo-sections and whole-mount zebrafish embryos^7^. MB has been used since the early 20^th^ century as a bright-field Nissl stain on thin sections. Here, we optimized the staining protocols for use with thick optically cleared human tissue samples requiring orders of magnitude lower concentrations than bright-field application.

The applied optimized labelling protocol was as follows: All samples were first incubated in freshly filtered solution of 50 % potassium disulfite (Sigma-Aldrich, 55777) in distilled water for 1h at RT. The samples were then washed for 1h at RT in distilled water. MG stock solution was prepared according to the method of Prieto et al.^7^. A 4 % aqueous MG (Sigma-Aldrich, 67060) solution was prepared and crystal violet impurities were removed by extractions with chloroform (Sigma-Aldrich, 372978), discarding the lower (violet) phase until no violet tinge could be observed in the lower phase. Stock solution was diluted 1:5000 in 0.1 M PBS with 1 % sodium azide (Sigma-Aldrich, S2002) at pH 7.4. Staining procedure for AO and NR was based on the protocols described in Schmued et al.^15^. For AO (CarlRoth, 7632.2) and NR (CarlRoth, T122.1) staining, a 1 % stock solution in 0.1 M PBS with 1 % sodium azide at pH 4 was prepared for each dye. 1 % MB (CarlRoth, A514.1) stock solution was prepared in PBS with 1 % sodium azide at pH 7.4. Samples for AO, NR and MB staining were incubated in a final concentration of 0.001 % in 0.1 M PBS with 1 % sodium azide at either pH 4 (AO, NR) or pH 7.4 (MB) for 1 day/mm tissue thickness (2-5 days total) at 4°C. All staining steps were carried out again in 6 well-plates in a volume of 6 ml/well on a shaker.

### Clearing and refractive index matching with MASH-RIMS

After labelling, samples were washed twice for 1 h in 0.1 M PBS of the respective pH and dehydrated in ascending concentration of methanol in water: 1 h each in 20 %, 40 %, 60 %, 80 %, and twice in 100 %. A volume of 5 ml per sample was used for each solution and incubation was performed at RT. Given the corrosiveness of the involved solutions, samples were then transferred into 50 ml incubation tubes made of high-density polyethylene (HDPE). Subsequently, samples were incubated for 3 h (2 mm thick samples) or overnight (5 mm thick samples; see **Supplementary Table 1**) in a mixture of 33 % methanol / 66 % dichloromethane (DCM, CarlRoth, 8424.2). Remaining methanol was washed out by incubation in 100 % DCM twice for 15 min (2 mm thickness) or twice for 1 h (5 mm). 50 ml tubes were filled completely with each solution.

For refractive index matching, cleared (dehydrated and delipidated) samples were incubated overnight at RT (transparency can already be achieved after several hours of incubation depending on sample thickness and the RIMS used) in 25 ml of one of the newly proposed MASH RIMS: 1) WGO/CA: 72 % methyl salicylate also known as wintergreen oil (WGO, Sigma-Aldrich, 84332) and 28 % *trans*-Cinnamaldehyde (CA, Sigma-Aldrich, C80687), 2) TDE/CA: 62 % *2,2*-Thiodiethanol (TDE, Sigma-Aldrich, 166782) and 38 % CA. The RIMS was changed once right before imaging and tubes were turned upside-down several times until no streaks were visible anymore in the fluid before mounting for microscopic imaging.

### Counterstaining cleared and MASH labelled samples with DAPI

Counterstains were performed with 4’,6-diamidino-2-phenylindole (DAPI) to label cell nuclei in cleared samples labelled with MASH-AO and MASH-NR. For DAPI labelling, 100 mg DAPI (CarlRoth, 6843.3) was dissolved in distilled water to prepare a 0.8 mg/ml stock solution. This stock solution was diluted 1:800 into the freshly prepared working solutions at the respective pH, resulting in a final concentration of 1 µg/ml. For labelling, each brain sample was incubated in a volume of 6 ml for 2 or 5 d at 4°C on a shaker for 2 or 5 mm thick samples respectively. For incubation 6 well-plates were used. For DAPI co-labelling of sections, an incubation time of 15 min at RT was used.

### MASH-RIMS property comparison

For refractive index matching of dehydrated and delipidated human tissue, the properties of several substances and solutions of various RI‘s were evaluated for their use as RIMS (**Supplementary Table 2**). As a previously unexplored substance for RIMS, *trans*-Cinnamaldehyde (CA, Sigma-Aldrich, C80687) was identified, an essential oil with a very high RI of 1.62 and low melting point which is mixable with TDE and WGO.

The following mixtures were evaluated for their clearing capacity of the dehydrated and delipidated adult human brain samples: 80 % glycerol in 0.1 M PBS (pH 7.4, RI = 1.44) and mineral oil (Sigma-Aldrich, M5904) with an RI of 1.47, pure TDE (Sigma-Aldrich, 166782) with an RI of 1.52, pure WGO (RI = 1.54, Sigma-Aldrich, 84332), and pure ethyl cinnamate (ECi, RI = 1.56, Sigma-Aldrich, 112372). Furthermore, mixtures of either 72 % WGO and 28 % CA or 62 % TDE and 38 % CA with an RI of 1.56 respectively and a solution of 38 % TDE and 62 % CA with an RI of approximately 1.58 were tested. As a control, 0.1 M PBS was used. To evaluate corrosiveness, the compatibility with selected plastic materials was checked. Several containers made from polystyrene, polypropylene, high-density polyethylene and tetrafluoroethylene were incubated for up to one week with the solutions described above.

The two MASH RIMS TDE/CA (62 % TDE and 38 % CA, RI=1.56) and WGO/CA (72 % WGO and 28 % CA, RI=1.56) were identified as having the combined desirable properties of an ideal (and adaptable) RI, low photobleaching, low corrosiveness and convenient storage at low temperature (low melting point). These solutions render archival human brain samples highly transparent with a slight remaining amber color (**Fig. 2 b,g; Supplementary Fig 5**) typical for many solvent-based clearing approaches. The transparency achieved with the recently described ECi^16^ is similar to the WGO/CA and TDE/CA RIMS (**Supplementary Fig. 3**). However ECi has a melting point of 6-8 °C and samples cannot be stored in the fridge once immersed in the liquid. Overall the Overall the TDE/CA RIMS was preferred, because it is even less corrosive for plastic equipment than WGO/CA and had better properties for long-term cold storage (low melting point) than ECi.

### Standard histological sectioning

For the optimization of the final staining protocol (described above) as well as for the validation experiments, standard histological sections of the same brain tissue were used. Therefore, manually cut blocks of human neocortical tissue were sectioned on a vibratome (VT1200 S, Leica Mikrosysteme Vertrieb GmbH, Wetzlar, Germany) into 50 µm thick sections.

### Optimization of staining conditions for MASH dye protocols

Staining conditions for MASH-AO and MASH-NR, in terms of pH and pre-treatment were optimized for maximum contrast (**Supplementary Figure 3 and 4**). Before staining, sections were incubated in either 50 % freshly filtered potassium disulfite solution for 15 min, dehydrated and delipidated in 70 %, 100 % and 70 % methanol for 5 min each or treated first with potassium disulfite and then methanol. Control sections were incubated in 0.1 M PBS for 30 min (**Supplementary Figure 3 and 4**). Sections were then washed for 5 min in distilled water and stained with AO or NR in the respective dye solutions described above for 15 min and washed in 0.1 M PBS of the respective pH for 5 min twice. After that sections were mounted in Kaiser‘s glycerol gelatine (CarlRoth, 6474.1).

### Validation of specificity of MASH dye protocols

Three validation experiments were performed on standard sections. 1) MASH dye protocols MASH-AO, MASH-NR and MASH-MB were compared in the same sections with standard bright-field (BF) stain Cresyl Violet. 2) MASH dye protocol MASH-MG was compared in the same sections with standard fluorescent stain DAPI. 3) MASH dye protocols MASH-NR, MASH-MB and MASH-MG were compared in the same sections with higher concentrations of the same dye imaged as a bright-field stain.

For experiment 1, standard histological sections were pretreated with potassium disulfite solution and stained with MASH dye protocols MASH-AO, MASH-NR and MASH-MB as described. Sections were then dehydrated in 50 %, 70 % and 100 % methanol for 5 min each, delipidated in 66 % DCM/33 % methanol for 15 min, and washed twice in 100 % DCM for 5 min. This was followed by rehydration in 100 %, 70 %, 50 % methanol and twice in 0.1 M PBS of the respective pH for 5 min. Sections were mounted in Kaiser‘s glycerol gelatine. For comparison of the MASH dye protocols with bright-field stains, coverslips of the MASH-labelled sections were removed by immersion in warm water after fluorescent imaging. Subsequently, sections were washed in warm distilled water for 5 min. and stained with cresyl violet acetate (AlfaAesar, J64318) by first dehydration in 70 % and 100 % ethanol for 5 min and rehydration in 70 % ethanol and distilled water for 2 min. This was followed by incubation in freshly filtered 50 % aqueous potassium disulfite solution for 15 min and two washes in distilled water for 5 min each. For staining a filtered cresyl violet solution of 1.5 % in water, 1 % acetic acid and 1 % 1 M sodium acetate was used. Samples were stained for 5 min and subsequently washed in acetate buffer for 2 min. Differentiation and dehydration was performed in 70 %, 96 % and 100 % ethanol. Finally samples were immersed in xylol for 5 min twice and mounted in entellan^®^ (**Supplementary Figure 1 a – i**). For experiment 2 on MASH-MG, the same procedures were followed, but sections were co-labelled with DAPI as described above for DAPI counterstaining (**Supplementary Figure 1 j – l).**

For experiment 3, sections fluorescently labelled with MASH-MB, MASH-MG and MASH-NR were stained with higher concentrations of the same dyes as a BF control. To this end, coverslips were removed and the sections were washed as described above. Sections were then stained in aqueous stock solutions of MB or MG, or in 0.1 % of NR in 0.1 M PBS at pH 4 for 5 min respectively. Hereafter, sections were washed 5 min in 0.1 M PBS with a pH of 7.4 (MB and MG) or pH 4(NR) and differentiated in 70 % and 100 % ethanol. After two 3 min incubations in xylol, sections were mounted in entellan^®^ and imaged (**Supplementary Figure 2**).

### Magnetic resonance imaging

Data acquisitions on the occipital lobe host sample were performed on a research 9.4T Siemens MAGNETOM scanner (Siemens Healthcare, Erlangen, Germany) according to the methods described in ^17^. A 3D multi-echo Gradient echo (GRE) sequence was used to acquire 200 µm isotropic data (Repetition time /Echo times: 45ms/ 7.86, 14, 24 and 34ms, flip angle (FA) = 28deg, bandwidth (BW) = 120 Hz/px, matrix dimensions = 400x400x416). Quantitative T_2_* estimation was performed by fitting a mono exponential decay model and a 3D surface reconstruction was created using Brain Voyager QX v2.8.

### Light-sheet and two-photon microscopic imaging

Before two-photon microscopic imaging (TPM), stained and cleared human brain samples were transferred to a glass petri dish, and a cover slip with water drop was placed on top of each sample. For TPM imaging experiments, a two-photon laser scanning microscope (Leica TCS SP5 MP, Leica Mikrosysteme Vertrieb GmbH, Wetzlar, Germany), equipped with a HCX APO L 20x/1.00W water immersion objective was used. Working distance of the objective was 2 mm and the excitation source was a 140 fs-pulsed Ti:sapphire laser (Chameleon Ultra II, Coherent Inc., Santa Clara, CA, USA), mode-locked at 800 nm. To avoid photobleaching and tissue damage, laser power was kept at 11% resulting in approx. 25-50 mW, at the sample surface. Images and image stacks were acquired with Leica Application Suite Advanced Fluorescence (Leica Microsystems). TPM Image acquisition settings are detailed in **Supplementary Table 3**.

Light sheet fluoresscence microscopic (LSFM) imaging was performed with the Ultramicroscope II (La Vision Biotech, Bielefeld, Germany), equipped with a SuperK Extreme Supercontinuum white light laser (EXW-12, NKT Photonics, Birkerød, Denmark). The used objective was a MVPLAPO 2X C/0,5 NA objective with dipping cap (Olympus, Japan), a working distance of 5.7 mm and a RI range of 1,33-1,56. The brain samples were fully immersed during the imaging process in 160 ml of imaging medium. TPM and LSFM acquisition settings for all experiments are detailed in **Supplementary Table 3**.

### Bright-field and fluorescence microscopy of thin tissue sections

For all imaging experiments on (non-cleared) vibratome sections an Olympus BX51WI DSU confocal microscope (Olympus, Center Valley, PA, USA) coupled to a Hamamatsu EM-CCD C9100 camera (Hamamatsu Photonics K. K., Hamamatsu, Japan) was used. The system was equipped with a motorized stage and a LEP MAC 5000 Controller System (Ludl electronic products, Hawthorne, NY, USA). Images were taken with either an Olympus PlanApo 2x/0.08 NA, 4x/0.16 NA or UPlanSApo 10x/0.40 NA objective (Olympus, Center Valley, PA, USA). Excitation and emission characteristics for all dyes are given in **Supplementary Table 2**. For acquisition the Stereo Investigator software (MBF Bioscience, Williston, Vermont, USA) was used.

### Microscopy data processing

For image post-processing such as brightness and contrast adjustments, subtraction of background, and thresholding, as well running depth-stack volume visualization, the open source software FIJI was used^18^. 3D Volume rendering was performed using Bitplane IMARIS. Cell body and tissue background signal analysis variation over imaging depth (**Figure 1h**) was performed in MATLAB using the Open Microscopy Environment (OME) MATLAB toolbox. At each depth of 500µm, 1500µm, 2500µm, 3500µm and 4500µm from the surface of the tissue, three consecutive data planes were used from the imaging stack for analysis (15 planes total). The middle 50% (in the light propagation direction) of each image plane was used to discard data away from the light sheet waist. At each of the five depths, a total of fifteen regions-of-interest (five per plane, three planes) of 3x3 pixels were selected for brightness assessment in both cell bodies and tissue background. The signal for each region-of-interest was taken as the average over the 3x3 pixel area. For both cell bodies and tissue background, boxplots were created depicting mean (’+‘ sign), median (box center line), 25th and 75th percentiles (box edges) and 9th and 91th percentiles (whiskers) of the n=15 region-of-interest signals at each depth.

## Data availability

The datasets generated and/or analyzed during the current study are available from the corresponding author on reasonable request.

## ACKNOWLEDGEMENTS

This work was supported by an ERC Starting Grant (MULTICONNECT, #639938 to AR, AS) and Dutch science foundation (NWO) VIDI Grant (#14637 to AR, SH). We thank Dr. Uwe Schroer (La Vision Biotech, Bielefeld, Germany) for assistance with light-sheet data acquisition. We thank Johannes Franz and Francisco Lagos for assistance with MRI and light-sheet data processing, and Hellen Steinbusch, Iulia Rusu and Annemarie Kiessling for assistance with tissue processing, labeling and lab work.

## AUTHOR CONTRIBUTION

SH identified protocol reagents and optimized protocols; SH, AS, RG & AR designed experiments; SH, AS & AH performed experiments; SH, AS & AR analyzed the data; SH, AS, AH, RG & AR wrote the paper.

## REFERENCES

1. Chung, K. et al. Structural and molecular interrogation of intact biological systems. Nature 497, 332–337 (2013).

2. Dodt, H.U. et al. Ultramicroscopy: three-dimensional visualization of neuronal networks in the whole mouse brain. Nature methods 4, 331–336 (2007).

3. Erturk, A. et al. Three-dimensional imaging of solvent-cleared organs using 3DISCO. Nature protocols 7, 1983–1995 (2012).

4. Kubota, S.I. et al. Whole-Body Profiling of Cancer Metastasis with Single-Cell Resolution. Cell reports 20, 236–250 (2017).

5. Renier, N. et al. iDISCO: a simple, rapid method to immunolabel large tissue samples for volume imaging. Cell 159, 896–910 (2014).

6. Susaki, E.A. et al. Whole-brain imaging with single-cell resolution using chemical cocktails and computational analysis. Cell 157, 726–739 (2014).

7. Prieto, D., Aparicio, G., Morande, P.E. & Zolessi, F.R. A fast, low cost, and highly efficient fluorescent DNA labeling method using methyl green. Histochemistry and cell biology 142, 335–345 (2014).

8. Schmued L. C., Swason L. W. & E., S.L. Some Fluorescent Counterstains for Neuroanatomical Studies. J Histochem Cytochem 30, 123–128 (1982).

9. Pan, C. et al. Shrinkage-mediated imaging of entire organs and organisms using uDISCO. Nature methods 13, 859 (2016).

10. Liebmann, T. et al. Three-Dimensional Study of Alzheimer’s Disease Hallmarks Using the iDISCO Clearing Method. Cell reports 16, 1138–1152 (2016).

11. von Economo, C. & Koskinas, G. Die Cytoarchitektonik der Hirnrinde des Erwachsenen Menschen: nTextband und Atlas mit 112 Mikrophotographischen Tafeln. (Springer, Vienna; 1925).

12. Kumar, A. et al. Dual-view plane illumination microscopy for rapid and spatially isotropic imaging. Nature protocols 9, 2555–2573 (2014).

13. Preibisch, S., Saalfeld, S., Schindelin, J. & Tomancak, P. Software for bead-based registration of selective plane illumination microscopy data. Nature methods 7, 418–419 (2010).

14. Renier, N. et al. Mapping of Brain Activity by Automated Volume Analysis of Immediate Early Genes. Cell 165, 1789–1802 (2016).

15. Schmued, L.C., Swanson, L.W. & Sawchenko, P.E. Some fluorescent counterstains for neuroanatomical studies. Journal of Histochemistry & Cytochemistry 30, 123–128 (1982).

16. Klingberg, A. et al. Fully Automated Evaluation of Total Glomerular Number and Capillary Tuft Size in Nephritic Kidneys Using Lightsheet Microscopy. Journal of the American Society of Nephrology: JASN 28, 452–459 (2017).

17. Sengupta, S. et al. High resolution anatomical and quantitative MRI of the entire human occipital lobe ex vivo at 9.4T. NeuroImage (2017).

18. Schindelin, J. et al. Fiji: an open-source platform for biological-image analysis. Nature methods 9, 676–682 (2012).

